# Whole genome sequence and comparative analysis of *Borrelia burgdorferi* MM1

**DOI:** 10.1101/200766

**Authors:** Neda Jabbari, Gustavo Glusman, Lena M. Joesch-Cohen, Panga Jaipal Reddy, Robert L. Moritz, Leroy Hood, Christopher G. Lausted

## Abstract

Lyme disease is caused by spirochaetes of the *Borrelia burgdorferi* sensu lato genospecies. Complete genome assemblies are available for fewer than ten strains of *Borrelia burgdorferi* sensu stricto, the primary cause of Lyme disease in North America. MM1 is a sensu stricto strain originally isolated in the midwestern United States. Aside from a small number of genes, the complete genome sequence of this strain has not been reported. Here we present the complete genome sequence of MM1 in relation to other sensu stricto strains and in terms of its Multi Locus Sequence Typing. Our results indicate that MM1 is a new sequence type which contains a conserved main chromosome and 15 plasmids. Our results include the first contiguous 28.5 kb assembly of lp28-8, a linear plasmid carrying the *vls* antigenic variation system, from a *Borrelia burgdorferi* sensu stricto strain.

## Introduction

Lyme disease is the most prevalent tick-borne disease in North America. Resulting from infections by spirochetes of the *Borrelia burgdorferi* sensu lato (s.l.) genospecies, the most frequent initial manifestation is erythema migrans, a characteristic expanding ring-shaped cutaneous lesion[1,2]. While some patients experience erythema migrans with or without flu-like symptoms, others may develop extracutaneous manifestations that can affect the nervous system (neuroborreliosis), the heart (Lyme carditis) or the joints (Lyme arthritis)[3–5]. What role pathogen genetic diversity plays in the producing varying symptoms is unknown. Currently, genomic sequence of forty seven strains of *Borrelia burgdorferi* sensu stricto (s.s.), the primary clinically relevant species in North America[6], are available from NIH GenBank. Of these, fewer than ten are described as complete assemblies. Given the complexity of Lyme manifestations and the scarcity of sequenced genomes, sequencing and characterization of more strains is vital for the understanding of *Borrelia* genetic diversity and disease manifestations.

The MM1 strain was isolated from the kidney of a white-footed mouse, *Peromyscus leucopus*, in Minnesota [7,8]. It was used in early studies of Lyme disease [7, 9–12] and it appears in patent applications for the development of *B. burgdorferi* detection assays and recombinant vaccines against Lyme disease [13,14]. However, its whole genome sequence has not yet been reported. We present here the full genome sequence of MM1, and provide comparative analysis, Multi Locus Sequence Typing (MLST), and annotation of its genome. Most notably, we report on the presence of the lp28-8 plasmid which contains its *vls* antigenic variation system.

## Materials and methods

### Growth conditions and DNA preparation

*B. burgdorferi* MM1 spirochetes were purchased from ATCC (ATCC 51990), resuspended and maintained in laboratory-produced media that resembles MKP/BSK-II media with 10% heat-inactivated rabbit serum (R7136-60, Sigma-Aldrich, USA), based on recipes provided by Dr. K. Strle[15,16]. ATCC 51990 MM1was first suspended in 10 mL of MKP/BSK-II media, then 1 mL of the diluted cells was inoculated into 13 mL of MKP/BSK-II media in 15 mL culture tubes (Corning tubes, USA) and incubated at 34°C, 5.0% CO_2_ with the lids of the tubes tightly sealed to minimize the oxygen level (microaerophilic condition). The seeded density of spirochetes was 5 ⨉ 10^6^ cells/mL and cells were grown until they reached mid-log phase (i.e., spirochete count reached 3 –5 ⨉ 10^7^). MM1 spirochetes four passages away from ATCC lot were harvested and washed with PBS buffer (pH 7.4) four times to remove the media components and pelleted, ready for subsequent DNA processing and genome sequencing.

Total DNA was isolated from pelleted MM1 spirochetes using DNeasy blood and tissue kit (Qiagen, USA). RNase A (Qiagen) was used to digest the RNA content as recommended by the manufacturer’s protocol. DNA was purified using the Agencourt AMPure XP method (Beckman Coulter, USA) and quantified with the Quant-iT PicoGreen dsDNA Assay Kit (ThermoFisher Scientific, USA) and a NanoDrop ND-1000 spectrophotometer. Integrity of the isolated DNA was checked on a 0.6 % agarose gel as well as with an Agilent DNA 12000 Kit (Agilent Technologies, Inc., USA).

### Genome sequencing

DNA sequence data were obtained primarily using the Pacific Biosciences (PacBio) Single Molecule, Real-Time (SMRT) system with DNA/Polymerase Binding Kit P6 v2, MagBead Kit v1 and DNA Sequencing Reagent Kit 4.0 v2 at the PacBio Sequencing Services of the University of Washington (Seattle, WA, USA). Long DNA reads were collected in one SMRT cell on a PacBio RSII instrument. Additional long reads were obtained using the Oxford Nanopore Technologies (ONT) MinION system. The MinION genomic DNA library was prepared according to the ONT Nanopore Sequencing Kit SQK-MAP006 protocol and quantified with Quant-iT PicoGreen dsDNA Assay Kit (ThermoFisher Scientific, USA). Sequencing reads were generated on Flow Cells (Flo-MAP003) with MinKNOW 0.51.1.62 and base-called with Metrichor 2.39. “2D pass” reads were converted into FASTA format with poretools v. 0.5.1. These reads are the highest quality reads, selected from reads where both strands of a double-stranded fragment were analyzed together in one pore.

### Genome assembly

*De novo* assembly of PacBio reads was performed with Hierarchical Genome Assembly Process (HGAP v2) using a minimum seed read length of 7800 bp. Contigs were merged or circularized with Circlator (v 1.5.0) [17]. In order to acquire the final consensus sequence, raw reads were filtered and mapped to the *B. burgdorferi* MM1 assembly with the resequencing protocol as implemented in SMRT Analysis Software v2.3.0.

### Sequence characterization and visualisation

Tandem repeats were defined with Unipro UGENE v1.9.8 as follows: 1–9 bp, 10–100 bp and >100 bp unit sizes were termed micro-, mini- and macro-satellite, respectively, and overlapping tandem repeats were excluded [18]. BLAST alignments of the sequences along with annotations were plotted using Easyfig (v 2.2.2) [19] and the Artemis Comparison Tool (ACT; Release 13.0.0)[20].

### *B. burgdorferi* MM1 plasmid identification and nomenclature

The plasmid content of MM1 was first evaluated visually with respect to the standard type strain B31. Reads were mapped to the B31 genome (assembly GCA_000008685.2) with the “resequencing” protocol included in PacBio’s SMRT Analysis software v2.3.0 (S1 Fig). Next, MM1 plasmid identities were confirmed by the sequence type of the *Borrelia* paralogous family (PFam) 32 protein encoded by each plasmid as defined by Casjens *et al*. [21]. Specifically, the protein sequences of MM1 and three fully-sequenced *B. burgdorferi* s.s. strains (B31, N40 and JD1; assemblies GCA_000008685.2, GCA_000166635.2 and GCA_000166655.2, respectively) were aligned with BLAST using an E-value cutoff of 0.001. These alignments were then clustered using Spectral Clustering of Protein Sequences (SCPS) [22] to infer homology relations between sequences based on pairwise similarity scores. Members of the four protein families, PFam32, PFam49, PFam50 and PFam57/62, as reasonable candidates for functioning in plasmid replication and partitioning [23], were identified on each contig (S1 Table). Plasmids were assigned names according to the PFam32 type matching their encoding PFam32 member (>95% amino acid sequence identity). Two exceptions were made to this method: cp9 and lp28-8. The smallest plasmid, cp9, does not encode PFam32, so it was identified by the sequence type of its PFam49, PFam50 and PFam57/62 members (S1 Table). Other than cp9, short contigs with no identifiable PFam32 type were not considered. The PFam32 protein from what would eventually be identified as the MM1 lp28-8 plasmid shared only a 60% amino acid sequence identity with PFam32 sequence type from B31 and JD1 cp32-3 plasmids. Instead, a protein BLAST against the NCBI non-redundant protein database, revealed it as fully identical to PFam32 type on *B. burgdorferi* 94a lp28-8 plasmid.

The MM1 lp54 plasmid subtype was identified following the criteria by Casjens *et al*.[24]. Dot plot analysis was performed to identify large (>400 bp) insertions, inversions or deletions in MM1 lp54 plasmid relative to lp54 subtypes. Nucleotide BLAST sequence comparisons and overall identical plasmid organization were investigated with Artemis Comparison Tool (ACT; Release 13.0.0) [20].

Correct replicon identification was confirmed visually by creating genome-scale dotplots of MM1 versus itself and versus B31 (S2 Fig) using GEPARD v1.4[25].

### Functional annotation

Genome features were annotated with RAST v2.0 [26] with the initial NCBI Taxonomy ID set to *B. burgdorferi* (139). To create a comparative annotation map of main chromosomes, FASTA sequences of all *B. burgdorferi* strains with complete genome assemblies were downloaded from NCBI GenBank. B31, CA382, JD1, N40, PAbe, PAli, and ZS7 (accession numbers AE000783.1, CP005925.1, CP002312.1, CP002228.1, CP019916.1, CP019844.1, and CP001205.1, respectively). The main chromosomes of B31, CA382, JD1, N40, and ZS7 were re-annotated with RAST annotation server using the same settings used for MM1 genome annotation. GenBank annotations for PAbe and PAli were used without modification.

### Phylogenetic analysis

A phylogenetic tree was constructed to evaluate the relationship of MM1 with respect to seven prominent laboratory *B. burgdorferi* s.s. strains. The tree was inferred from the whole main chromosome sequence data of s.s. strains MM1, B31, CA382, JD1, N40, PAbe, PAli, and ZS7 using REALPHY v.1.12[27] with outgroup *B. bissettii* DN127 (CP002746.1). The strains were ordered for plotting based on genomic distance from MM1 as determined by GGDC v.2.1[28] with its default settings for formula two. The tree was plotted using TreeGraph2 v2.13[29].

The eight chromosomal housekeeping genes *clpA*, *clpX*, *nifS*, *pepX*, *pyrG*, *recG*, *rplB* and *uvrA* of the Multi Locus Sequence Typing (MLST) scheme [30] were used to determine MM1 Sequence Type (ST). Corresponding locus sequences in MM1 were queried against the *Borrelia* MLST database [31]. Alleles with perfect matches to the database were assigned type numbers. Where perfect matches did not exist, single-base mismatches were noted and confirmed using Oxford Nanopore data. For confirmation, the novel alleles were searched against all MinION filtered 2D reads using FASTA v.36[32]. Hits were aligned with Clustal Omega v.1.2.1[33] and visually inspected (S3 Fig).

## Results and discussion

### Overall description of the *B. burgdorferi* MM1 genome

The MM1 genome, comprising a single large chromosome and fifteen plasmids, was determined to be 1,280,240 bp in size based on an average coverage depth of 737× that ranged from 351× to 2312× across the replicons (Table 1). The main linear chromosome represents 71% of the genome size (908,512 bp), with the plasmid content providing the remaining 29% (371,728 bp). According to RAST analysis, the MM1 genome carries 1338 annotated genes with 865 (65%) on the main chromosome including 36 RNA coding genes and 473 plasmid features (Table 1). Of all the annotated features, 37% are connected to a subsystem, the components of which are mainly (92%) on the chromosome (Table 1).

**Table 1.**
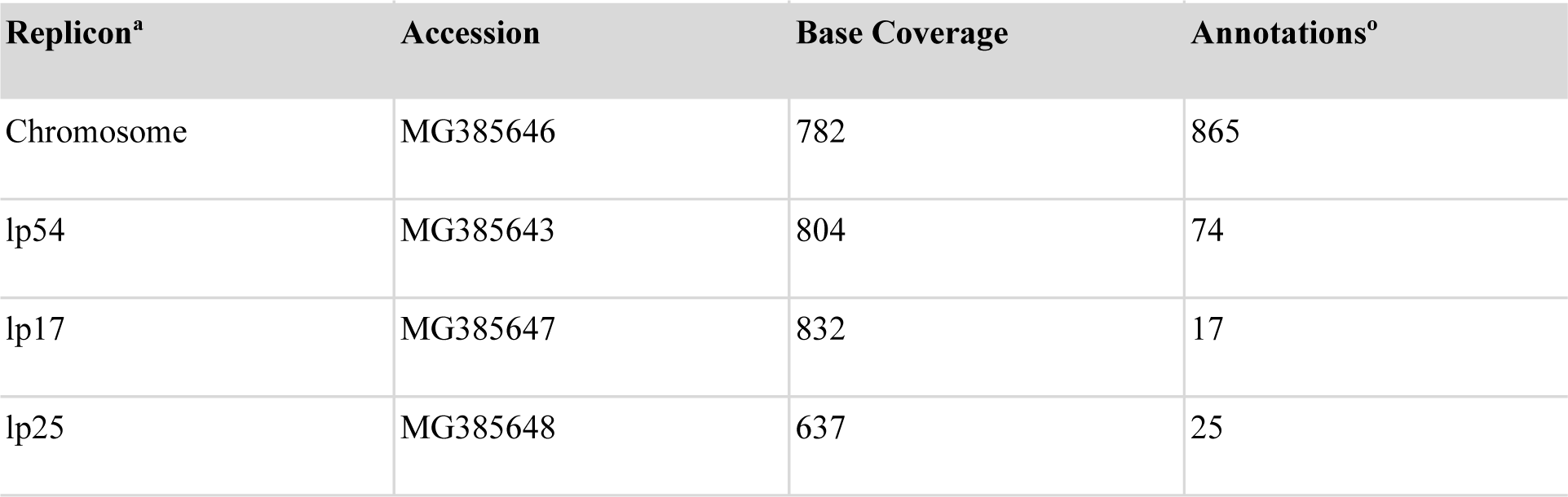
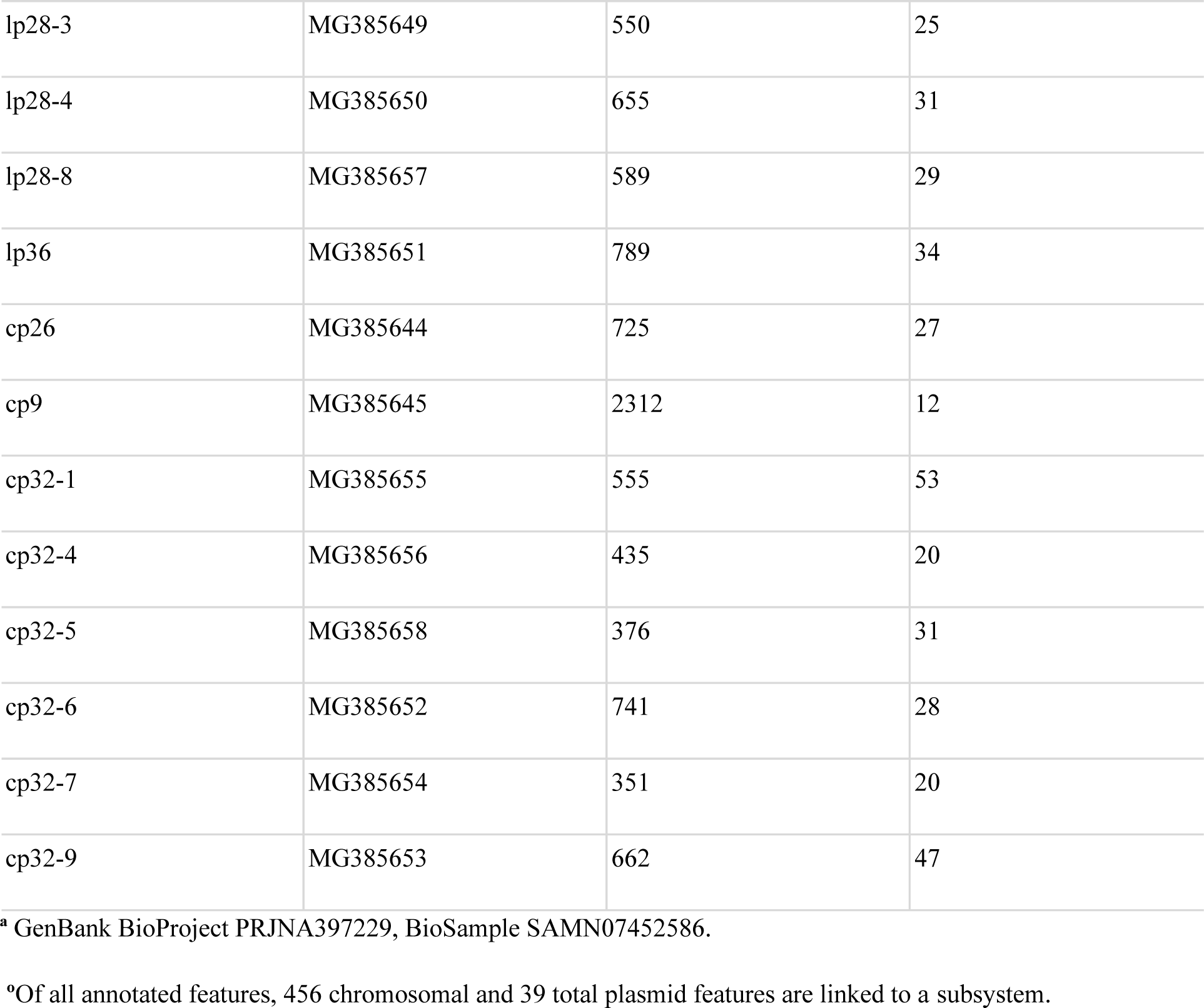
**GenBank, Coverage and Annotation Details for *B. burgdorferi* MM1 Genome**

From among the subsystem categories present in the MM1 genome, Protein Metabolism and Sulfur Metabolism have the highest and lowest subsystem feature counts, respectively. Interestingly, all the subsystem categories except for “Phages, Prophages, Transposable elements, Plasmids” category are also present in the s.s. B31 genome (Table 2).

**Table 2.** **Subsystem Categories Present in MM1 Genome in Relation to B31**

| **Category** | **Subsystem feature counts** |  |
| --- | --- | --- |
|  | **MM1** | **B31** |
| Protein Metabolism | 171 | 164 |
| RNA Metabolism | 68 | 82 |
| Motility and Chemotaxis | 62 | 58 |
| DNA Metabolism | 55 | 55 |
| Carbohydrates | 48 | 49 |
| Nucleosides and Nucleotides | 38 | 34 |
| Fatty Acids, Lipids, and Isoprenoids | 37 | 35 |
| Cell Wall and Capsule | 36 | 41 |
| Amino Acids and Derivatives | 32 | 32 |
| Cell Division and Cell Cycle | 26 | 26 |
| Virulence, Disease and Defense | 22 | 23 |
| Stress Response | 21 | 15 |
| Membrane Transport | 18 | 15 |
| Cofactors, Vitamins, Prosthetic Groups, Pigments | 17 | 21 |
| Phosphorus Metabolism | 10 | 14 |
| Respiration | 9 | 9 |
| Regulation and Cell signaling | 7 | 12 |
| Potassium metabolism | 6 | 4 |
| Phages, Prophages, Transposable elements, Plasmids | 6 | – |
| Miscellaneous | 3 | 10 |
| Dormancy and Sporulation | 2 | 2 |
| Sulfur Metabolism | 1 | 1 |

Given that the plasmid profile of *B. burgdorferi* may undergo changes as a result of prolonged *in vitro* cultivation [34–36], our sequenced low-passage culture of MM1 (see Materials and methods), like other sequenced *Borrelia*, may be no longer homogeneous with some plasmids lost during culturing.

### The main, linear chromosome of *B. burgdorferi* MM1

*B. burgdorferi* MM1 carries a main linear chromosome of 908,512 bps in length with a GC content of 28.6%—similar in size (902,191 - 922,801 bp) and GC content (28.5 - 28.6%) to the main chromosomes of other s.s. strains that have a complete genome assembly (Table 3). Consistently, the MM1 chromosome shares a 99% nucleotide sequence identity and a similar tandem repeat profile (e.g., microsatellite, minisatellite and macrosatellite) with these other main chromosomes (Table 3).

**Table 3.**
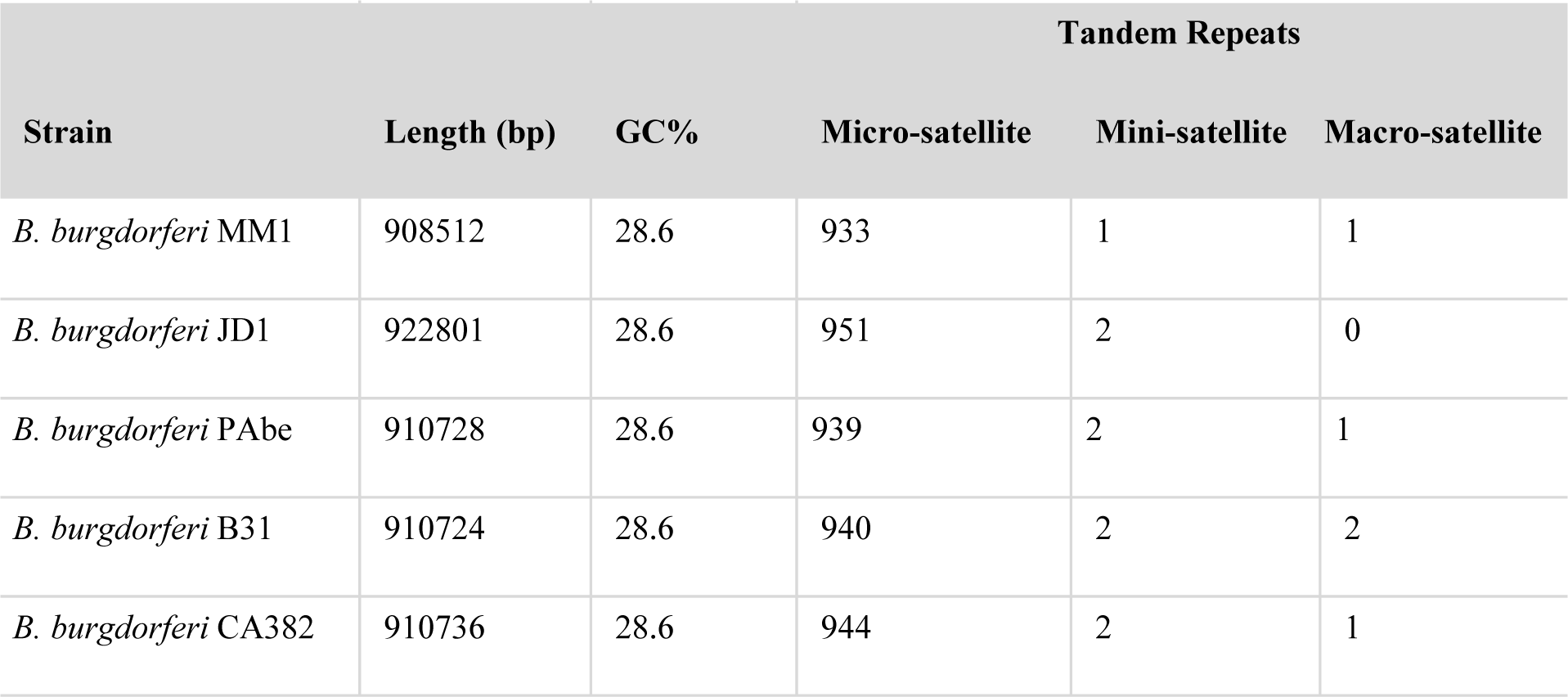
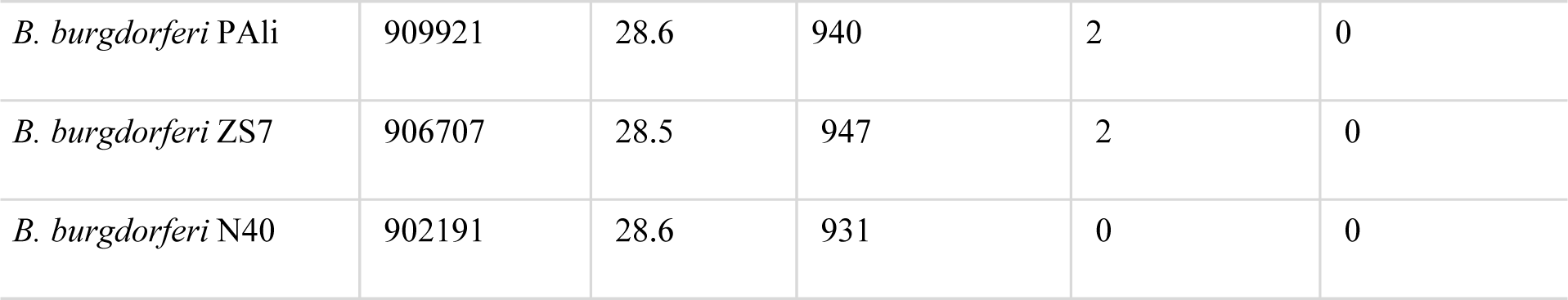
***B. burgdorferi* MM1 Chromosome in Relation to Representative *B. burgdorferi* Strains**

With the exception of the right end extension, the chromosomal gene contents of sequenced s.s. strains are essentially identical [37,38]. Accordingly, comparative annotation map of MM1 and s.s. strains suggests a consistent overall gene synteny over the entire chromosome with some variation in the right end extension (Fig 1).

**Fig 1.**
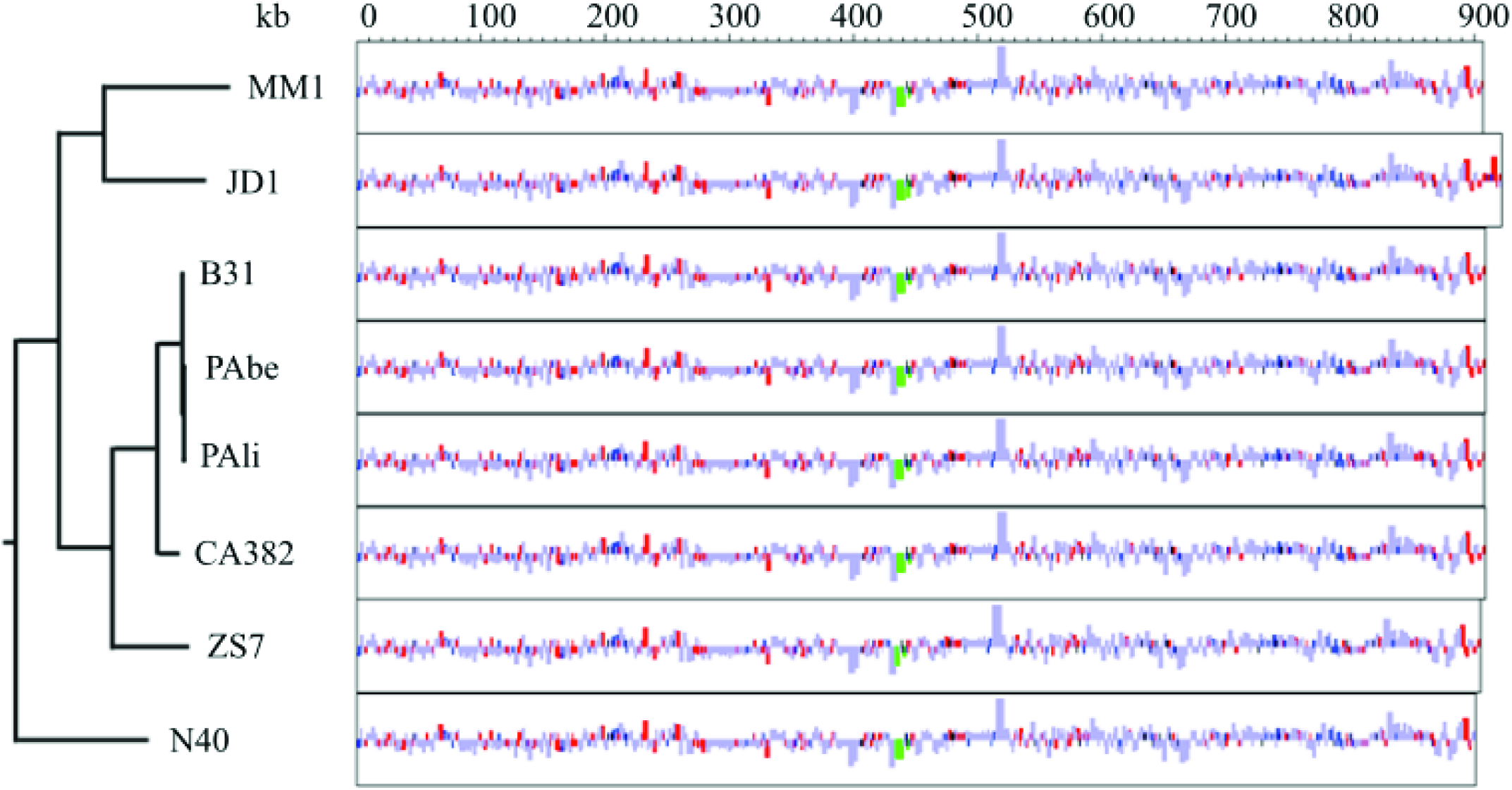
Comparative annotation map of the main chromosome of *B. burgdorferi* s.s. strains with complete genome assemblies. Rectangles represent annotated genes, color-coded by type: CDS in light blue, predicted CDS in dark blue, hypothetical CDS in red, rRNA in green and tRNA in black. Rectangle heights denote gene length; genes shorter than 1 kb are displayed enlarged for visibility. Features above or below the midline for each strain represent genes transcribed on the top or bottom strand, respectively. The phylogenetic tree was inferred from the whole main chromosome sequence data and rooted using *B. bissettii* DN127 as outgroup.

### *B. burgdorferi* MM1 plasmid content and variation from *B. burgdorferi* strains

An initial mapping of raw sequence reads from the MM1 genome to the reference strain B31 (see Materials and Methods) indicated less than 26% base calling for B31 linear plasmids (lp) lp21, lp28-1, lp28-2, lp38, lp56 and lp5 suggesting their absence in MM1 genome. However, the presence/absence of other B31 plasmids and, in particular, the circular plasmid (cp) cp32s that are homologous nearly throughout their lengths in *B. burgdorferi* [37] remained ambiguous. The observed depth of coverage across B31 plasmids is depicted in S1 Fig.

According to the PFam32 nomenclature scheme [21], the sequenced culture of MM1 carries fifteen plasmids encoding seven linear and seven circular plasmid PFam32 types. The small cp9 plasmid that does not encode a PFam32 protein defines an additional plasmid compatibility type in MM1 genome. While some plasmids might be spontaneously lost between original isolation and the low passage culture used for sequencing, the number of identified plasmids in MM1 is within the previously defined range in a set of 14 *B. burgdorferi* strains (e.g. 6-11 and 6-12 circular and linear sequenced plasmids, respectively) [24].

A comparison of variation in plasmid profile, size and GC content among MM1, B31, JD1 and N40 strains, is shown in Table 4. Similar to the type strain B31, MM1 carries linear plasmids lp54, lp17, lp25, lp28-3, lp28-4, lp36 and circular plasmids cp9, cp26, cp32-1, cp32-4, cp32-6, cp32-7, cp32-9. Among the non-cp32 plasmids, lp17, lp25, and lp36 appear to have major structural differences between the two strains (Table 4, Fig 2). The MM1 genome carries the linear plasmid PFam32 type, lp28-8, which is not present in the sequenced B31. In addition, MM1 carries cp32-5 PFam32 type, which is not encoded by B31 plasmids, but is encoded by the circular plasmids of N40 and present in strains JD1 in fusion with cp32-5 as cp32-1+5 (Table 4) [37].

**Table 4.** ***B. burgdorferi* MM1 plasmid content in relation to *B. burgdorferi* B31, JD1 and N40 strains.**

| Replicon<sup>a</sup> | Code | MM1 |  | B31 |  | JD1 |  | N40 |  |
| --- | --- | --- | --- | --- | --- | --- | --- | --- | --- |
|  |  | Length | GC% | Length | GC% | Length | GC% | Length | GC% |
| lp54 | A | 53.8 | 28.1 | 53.7 | 28.1 | 52.9 | 28.2 | 53.8 | 28.1 |
| lp17 | D | 14.4 | 23.7 | 16.8 | 23.1 | 17.5 | 23.9 | 20.7 | 24.3 |
| lp21 | U | – | – | 18.8 | 20.7 | – | – | – | – |
| lp25 | E | 24.2 | 25.4 | 24.2 | 23.4 | 22.8 | 25.4 | 23.4 | 23.3 |
| lp28-1 | F | – | – | 28.2 | 32.8 | 24.4 | 32.7 | – | – |
| lp28-2 | G | – | – | 29.8 | 31.6 | – | – | 29.5 | 31.7 |
| lp28-3 | H | 26.8 | 25.0 | 28.6 | 25.0 | 28.8 | 25.0 | – | – |
| lp28-4 | I | 27.5 | 24.5 | 27.3 | 24.5 | 30.1 | 24.7 | 29.4 | 24.6 |
| lp28-5 | Y | – | – | – | – | 24.6 | 23.9 | 26.6 | 25.3 |
| lp28-6 | Z | – | – | – | – | 27.0 | 32.4 | – | – |
| lp28-7 | AA | – | – | – | – | 30.0 | 31.6 | – | – |
| lp28-8° | RC | 28.5 | 33.2 | – | – | – | – | – | – |
| lp36 | K | 28.0 | 25.5 | 36.8 | 26.9 | 22.8 | 25.6 | 29.2 | 31.6 |
| lp38 | J | – | – | 38.8 | 26.1 | 27.6 | 22.9 | 37.9 | 26.0 |
| lp5 | T | – | – | 5.2 | 23.8 | – | – | – | – |
| lp56 | Q | – | – | 53.0 | 27.3 | – | – | – | – |
| cp26 | B | 26.5 | 26.3 | 26.5 | 26.3 | 26.5 | 26.2 | 26.5 | 26.3 |
| cp9 | C | 11.4 | 24.3 | 9.4 | 23.7 | – | – | 8.7 | 24.3 |
| cp32-1 | P | 30.9 | 29.3 | 30.8 | 29.4 | 60.7<sup>n</sup> | 29.2 | – | – |
| cp32-3 | S | – | – | 30.2 | 28.9 | 29.7 | 28.9 | – | – |
| cp32-4 | R | 13.3 | 26.8 | 30.3 | 29.3 | – | – | 15.9 | 27.0 |
| cp32-5 | V | 21.9 | 28.5 | – | – | 60.7<sup>n</sup> | 29.2 | 29.6 | 29.2 |
| cp32-6 | M | 20.0 | 27.7 | 29.8 | 29.3 | 30.8 | 29.0 | – | – |
| cp32-7 | O | 14.0 | 26.3 | 30.8 | 29.1 | – | – | 17.5 | 27.2 |
| cp32-8 | L | – | – | 30.9 | 29.1 | 31.1 | 29.1 | – | – |
| cp32-9 | N | 30.6 | 29.1 | 30.7 | 29.3 | 30.7 | 29.2 | 30.7 | 29.0 |
| cp32-10<sup>y</sup> | Q | – | – | – | – | 30.3 | 29.3 | 30.2 | 29.2 |
| cp32-11 | W | – | – | – | – | 29.9 | 29.0 | – | – |
| cp32-12 | X | – | – | – | – | 30.0 | 29.3 | 27.8 | 28.1 |
| **Total plasmids** |  | 15 : 7 (L), 8 (C) |  | 21: 12 (L), 9 (C) |  | 20: 11 (L), 9 (C) |  | 16: 8 (L), 8 (C) |  |

<sup>a</sup>Length is listed in kbps.
<sup>o</sup>In strains of *B. afzelii*, where lp28-8 is documented (e.g. K78 and PKo), code “AC” is initially given to plasmid lp28-8 and later revised to “RS”, whereas, in others (e.g. *B. burgdorferi* strain 94a, *B. valaisiana* VS116, *B. spielmanii* A14S), code N is initially given and later revised to “RS” [24]. Consistently, we designate “RS” as the code for MM1 lp28-8 plasmid.
<sup>n</sup>JD1cp32-1 and cp32-5 are described as fused into 60.7 kbps circular cp32-1+5 plasmid [37]. <sup>y</sup>In B31 cp32-10 is integrated into lp56 plasmid [37]. L, linear; C, Circular.

**Fig 2.**
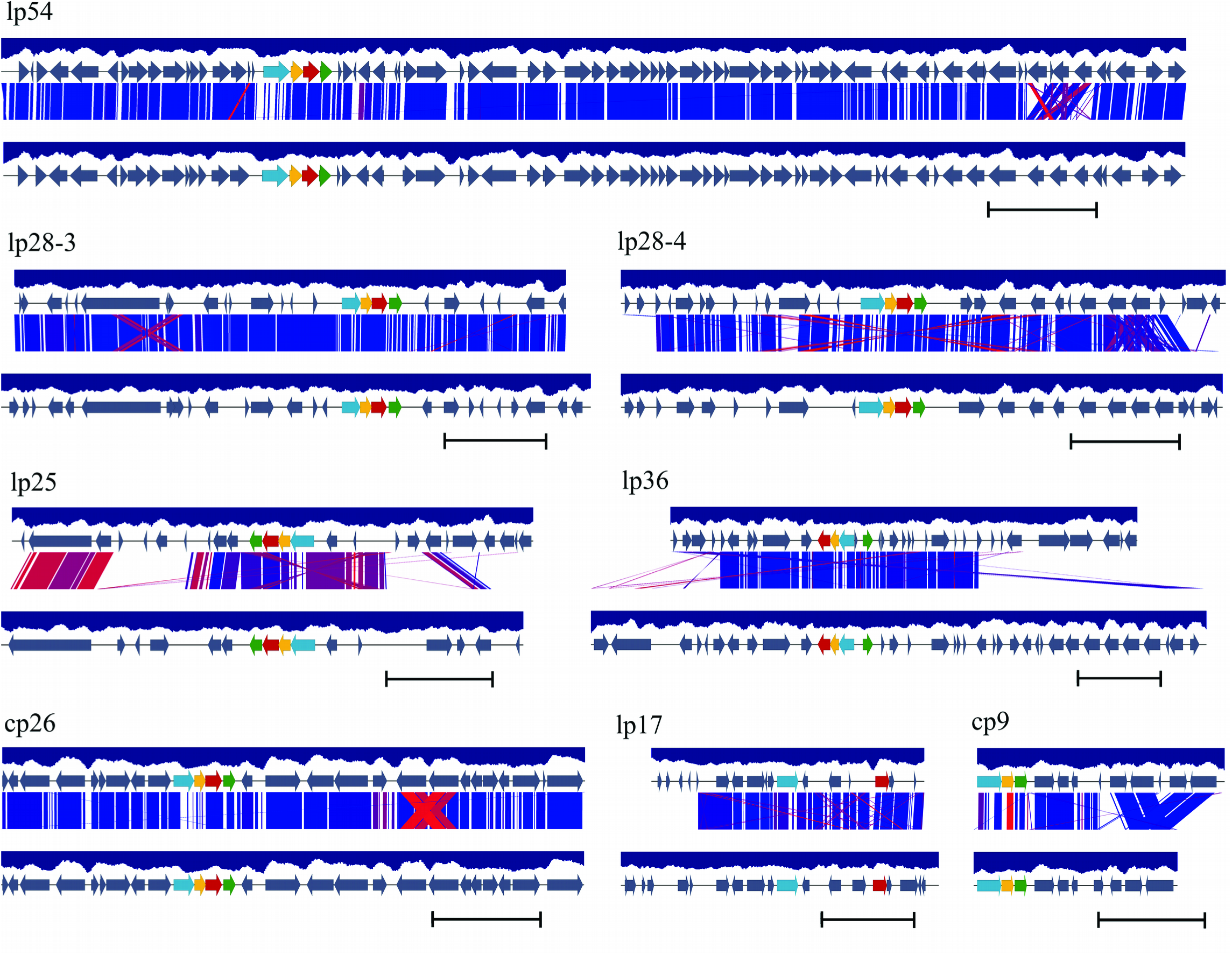
A diagrammatic depiction of the differences in the organization of the non-cp32 plasmids present in *B. burgdorferi* MM1 and B31 strains. Annotated genes with or without CDSs are represented as grey arrows. PFam organization and synteny is displayed on each plasmid in MM1 (top) and B31 (bottom) strains as red (PFam32), green (PFam49), yellow (PFam50) and blue (PFam57/62) arrows. Among B31the 4 PFams in B31, genes that appear to be disrupted and pseudogene relatives are not color coded [23] and only PFam types with consecutive numbers in MM1 are color coded. BLAST alignments between plasmid pairs (E-value <0.001) are indicated by blocks ranging in color from red (80% sequence identity) to dark blue (100% sequence identity). GC content is calculated in a window of 400 bps and displayed as a blue histogram bar of a GC content lower than 50%. Sequences are shown in full and drawn to scale with scale bars representing 5 kbps.

### The plasmids lp54 and cp26 in *B. burgdorferi* MM1

The linear plasmid lp54 and the circular plasmid cp26 have for the most part conserved synteny among the s.l. genomes [39,40]. Among the lp54 plasmid genes with low sequence identity across *Borrelia* species, are the genes encoding decorin binding proteins A (DbpA) and B (DbpB) that are critical for the overall virulence of *B. burgdorferi [41]*. DbpA is highly heterogeneous between and within s.l. species with as low as 70% sequence identity within s.s. strains, whereas DbpB is more conserved (>96% identity) within s.s. [42–44]. Consistent with the substantial difference in their sequence heterogeneity among *Borrelia* isolates, DbpA and DbpB proteins of MM1 (corresponding gene locus tags BbuMM1_A250 and BbuMM1_A260, respectively) share 79% and 100% sequence identity to B31 (Fig 3).

**Fig 3.**
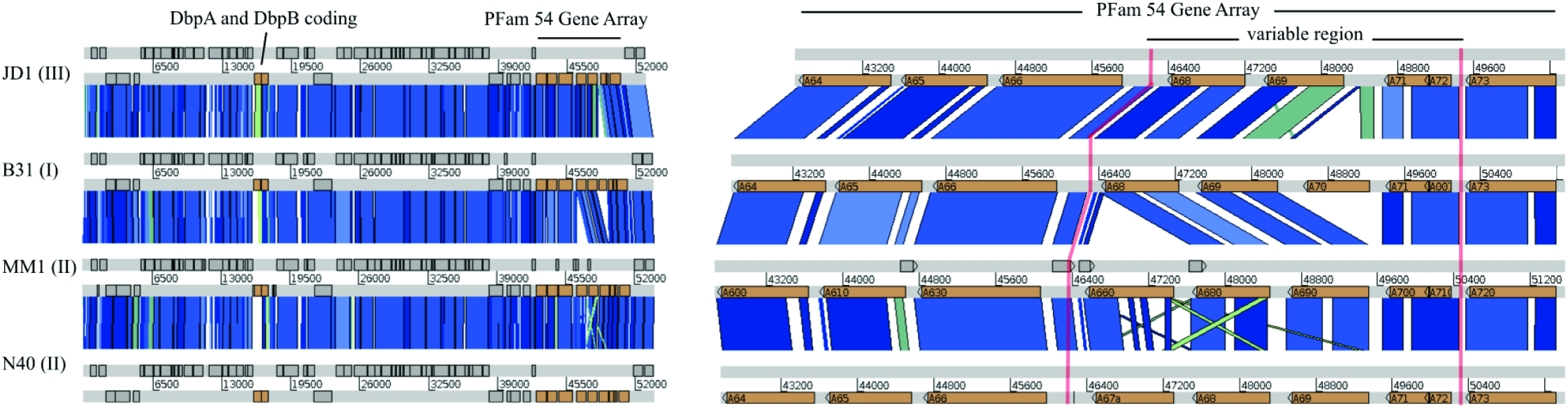
Synteny map of lp54 containing the PFam54 gene array in MM1, JD1, B31 and N40 strains. Left: The lp54 plasmid of MM1 *B. burgdorferi* in comparison to JD1, B31 and N40 strains.The strains are indicated in the left side of the map along with organizational subtypes indicated by Roman numerals in parenthesis. Plasmid is represented by a horizontal white bar labeled with positions with annotations on light grey bars. BLAST alignments between the lp54 assemblies are indicated by blocks ranging in color from green (95% sequence identity) to dark blue (100% sequence identity) with a minimum Score Cutoff of 28. Annotated features are indicated in dark grey and orange (dbpA and dbpB genes, PFam54 gene array). Right: A zoomed view of PFam54 gene array of lp54 plasmid in JD1, B31, MM1 and N40 strains. PFam54 variable region is sandwiched by *bba64*, *bba65*, and *bba66* on one side, and *bba73* on the other side. Among the 4 strains, MM1 variable region is highly conserved with N40.

Near one terminus of the lp54 plasmid, we identified a PFam54 gene array of close homologs of complement regulator-acquiring surface protein 1, CRASP-1, that has the most variable gene content caused by gene duplication, loss and sequence diversification [45,46]. This variable region is between the genes homologous to *bba66* and *bba73* in a set of ten *Borrelia* lp54 plasmids [45]. The gene order within MM1 lp54 variable region is very different from that in B31; MM1 carries Bbu_MM1A660 located between genes homologous to *bba66* and *bba68*, whereas B31 carries the *bba70* gene that is absent in MM1 (Fig 3). A comparison of the lp54 plasmid variable region in MM1 and N40 strains reveals highly conserved synteny and sequence, with 99% sequence identity between the Bbu_MM1A660 and BbuN40_A67a genes (Fig 3). MM1 and N40 lp54 plasmids have conserved gene order throughout their length. They do not carry insertions, inversions or deletions larger than 400 bps and inter-plasmid DNA exchanges relative to one another [24], therefore, MM1 lp54 plasmid appears to be a type II lp54.

*B. burgdorferi* cp26 plasmid is present in all natural isolates and encodes functions critical to bacterial viability [47]. The highly diverse outer-surface protein C (OspC) against which mammals develop protective immunity is encoded by the *ospC* gene on the cp26 plasmid [48]. MM1 strain carries a type U *ospC* allele with 100% nucleotide sequence identity to type U *ospC* alleles in northern US s.s. strains 94a and CS5[49] 84% sequence identity with type A *ospC* allele in B31 strain.

### *B. burgdorferi* MM1 plasmid lp28-8

MM1 plasmid lp28-8 encoding PFam32 protein is 100% and 90% identical in amino acid sequence to the lp28-8 encoding PFam32 proteins in *B. burgdorferi* 94a and *B. valaisiana* VS116, respectively. MM1 lp28-8 carries a typical cluster of PFam32, PFam49, PFam50, PFam57/62 that is 100% identical to the corresponding region of lp28-8 in *B. burgdorferi* 94a [24] and 86% identical to the corresponding region of lp28-8 in *B. afzelii* strains PKo and K78, *B. valaisiana* VS116 and *B. spielmanii* A14S (Fig 4).

**Fig 4.**
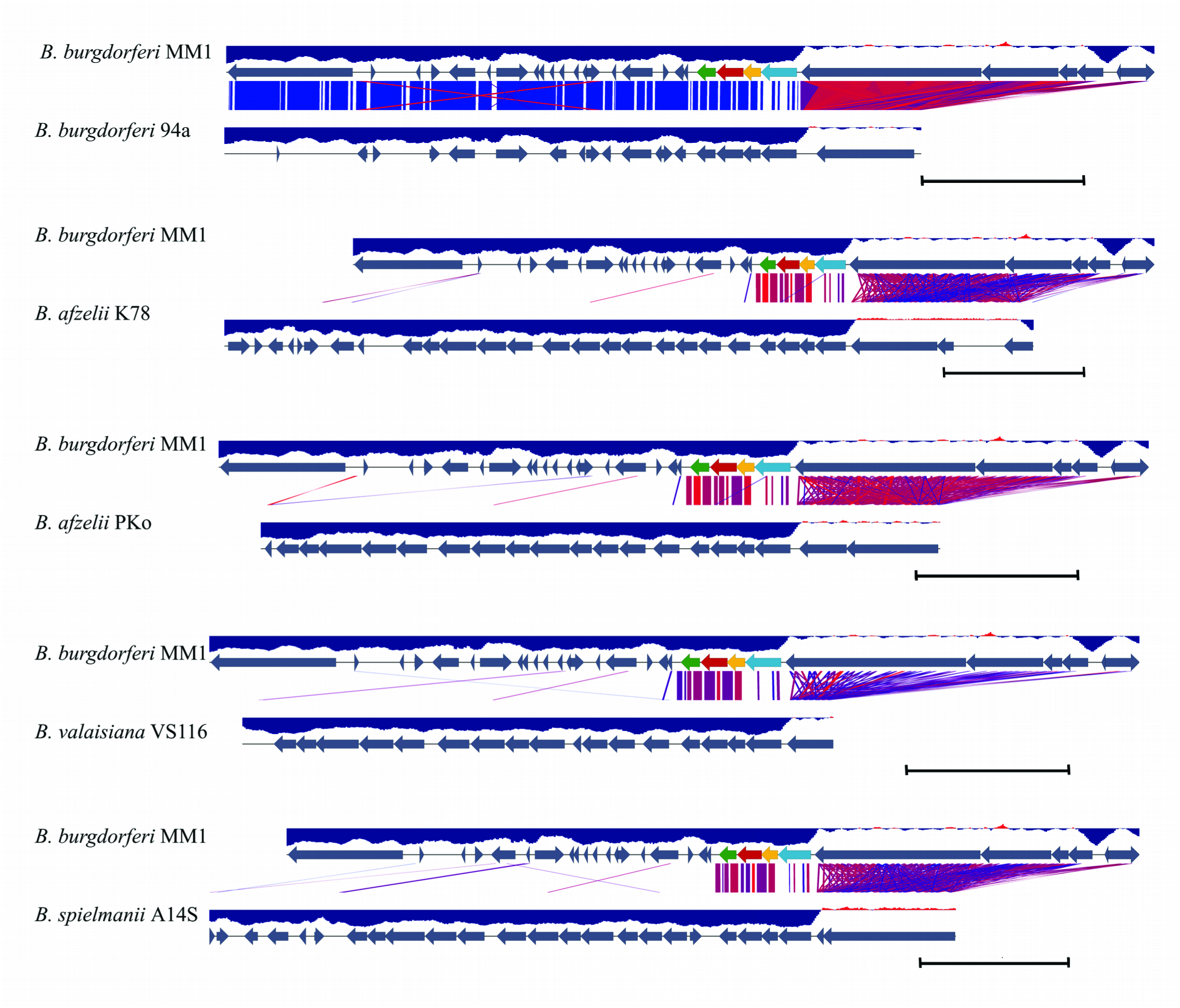
A diagrammatic depiction of the differences in the organization of the lp28-8 plasmids in MM1 and *B. burgdorferi* s.l. species. Annotations are represented as grey arrows. MM1 PFam organization and synteny are displayed on each alignment as red (PFam32), green (PFam49), yellow (PFam50) and blue (PFam57/62) arrows. Only PFam types with consecutive numbers are color coded. BLAST sequence comparison (E-value <0.001) represents a similarity range from 80% (red) to 100% (blue) among plasmid pairs. GC content is calculated in a window of 400 bps and displayed as a blue histogram bar of a GC content less or higher than 50%. The size of lp28-8 in *B. burgdorferi* s.s. 94a (largest contig), *B. afzelii* strain 78, *B. afzelii* strain PKo, *B. valaisiana* VS 116, and *B. spielmanii* A14S is 21,295, 28,638, 20715, 18022, and 24828 bps respectively. Sequences are shown in full and scale bars represent 5 kbps.

This assembly of the MM1 lp28-8 plasmid is the most complete for a *B. burgdorferi* s.s. to date. Previous contigs had not approached 28 kb [24,37,50]. In fact, it was recently described as a new PFam32 type present only in one s.s. strain, 94a, from among a set of 14 characterized s.s. isolates. Due to difficulties in assembling long repetitive tracts, the sequence of lp28-8 plasmid of 94a remains unfinished and reported as contigs of 21,295, 4,910 and 393 base pairs [24]. Our sequenced *B. burgdorferi* MM1 lp28-8 is a continuous contig of 28,515 bps in length that is 99% identical to lp28-8 in *B. burgdorferi* 94a over 95% of its sequence, and its leftmost 13 kb shares a 99% identity to plasmid lp38 subtype V. A comparison of sequence and organization of lp28-8 in MM1 to lp28-8 in *B. burgdorferi* 94a and s.l. species such as *B. afzelii* [51,52], *B. valaisiana* and *B. spielmanii* [53] is depicted in Fig 4.

MM1 lp28-8 plasmid carries a *vls* locus in one end with a characteristic high GC content [54]. The *vls* locus is required for long-term survival of Lyme *Borrelia* in infected mammals and is characterized with a *cis* location of the expression site *vlsE* and a contiguous array of *vls* silent cassettes [54]. Consistently, the average GC content of annotations within the *vls* locus of MM1 is 47% leading to an elevated average GC content of 33.2% for plasmid lp28-8, while the GC content range for the rest of MM1 genome is 23.7% to 29.3% (Table 4, Fig 4). MM1 lp28-8 includes an array of 18 sequences with at least 85% sequence identity to *vlsE* (Fig 5).

**Fig 5.**
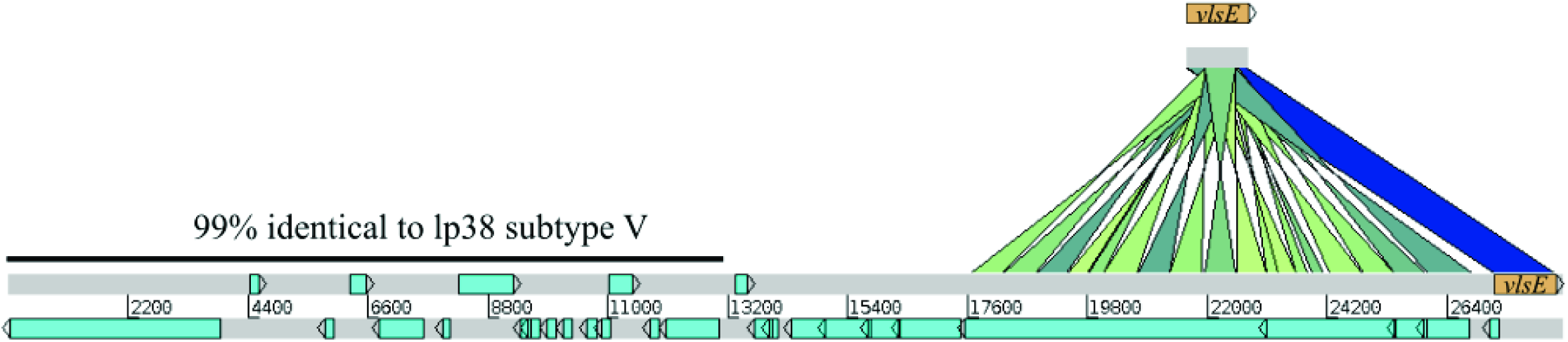
The *Vls* antigenic variation system in *B. burgdorferi* MM1. This locus on lp28-8 plasmid contains the expression site *vlsE* as well as a contiguous array of 18 DNA sequence blocks that are each at least 85% identical to *vlsE* (BbuMM1_RS290). BLAST alignments between the MM1 *vlsE* locus and lp28-8 plasmid are indicated by blocks ranging in color from green (85% sequence identity) to dark blue (100% sequence identity) with a minimum Score Cutoff of 85. The leftmost 13134 bps of lp28-8 plasmid is 99% identical to lp38 subtype V.

### The cp32 plasmids of *B. burgdorferi* MM1

The cp32 plasmid family members are described to have 12 PFam32 types in a set of 14 fully analyzed s.s. genome sequences [24]. The *B. burgdorferi* MM1 genome contains six members of the cp32 family, cp32-1, cp32-4, cp32-5, cp32-6, cp32-7, and cp32-9 (Table 4). Based on Spectral Clustering of Protein Sequence analysis, the MM1 cp32 plasmids consistently share an amino acid sequence identity of more than 99% with their PFam32 genes in either B31 or N40 strains (S1 Table). Among the six, cp32-5 plasmid is absent in *B. burgdorferi* B31.

### MLST reveals *B. burgdorferi* MM1 strain is a new sequence type

Based on the MLST scheme, no isolate in the *Borrelia* MLST database [31] was found to have the same allelic profile as MM1, therefore, we conclude that MM1 is a unique sequence type (ST). Of the eight loci used in the MLST scheme, six loci (*clpA*, *clpX*, *pyrG*, *recG*, *rplB* and *uvrA* excluding *nifS* and *pepX*) in the MM1 genome were the same allelic type as the corresponding loci in ST37 and ST18 s.s. isolates within the *Borrelia* MLST database (Table 5). *NifS* and *pepX* loci were partially matched (different in only 1 nt) to allelic types 12 and 1, respectively. The closest allelic profile to MM1 was observed in ST37 isolates with 1 bp mismatch in *nifS* loci. Interestingly, ST37 and ST18 isolates belong to Northeastern US or Eastern/East-central Canada, whereas MM1 is a Midwestern USA isolate.

**Table 5.** **MM1 MLST in relation to isolates (ST 37 and ST 18) with 6 out of 8 allelic types identical to MM1**

| isolate | country<sup>a</sup> | source<sup>o</sup> | allelic type |  |  |  |  |  |  |  |
| --- | --- | --- | --- | --- | --- | --- | --- | --- | --- | --- |
|  |  |  | *clpA* | *clpX* | *nifS* | *pepX* | *pyrG* | *recG* | *rplB* | *uvrA* |
| MM1<sup>o</sup> | US | M | 7 | 6 | – | – | 1 | 5 | 5 | 5 |
| ST37 | US (19), CA (2) | H (2), T(19) | 7 | 6 | 12 | 1 | 1 | 5 | 5 | 5 |
| ST18 | US (20), CA (5) | H (7), T (18) | 7 | 6 | 6 | 1 | 1 | 5 | 5 | 5 |

<sup>a</sup>All US locations are Northeastern US and Eastern/East-central Canada locations respectively. M, mouse; H, human, T, tick. <sup>o</sup>MM1 *nifS* and *pepX* loci match partially (1 nt difference) to *nifS*-12 and *pepX*-1, respectively.

## Conclusions

We report the full genome sequence and organization of *B. burgdorferi* MM1 derived from *in vitro* cultured spirochetes. Like other *B. burgdorferi* s.s. strains, MM1 appears to have a quite evolutionarily stable chromosome with little variation in size and content [21,37,38]. Our MM1 genome assembly produced 15 plasmids, of which seven are linear and eight are circular types. Based on the unique allelic profile of MM1 in MLST analysis, our study identifies MM1 as a new ST among Northern USA *B. burgdorferi* s.s. isolates. Except for the 94a strain, the lp28-8 plasmid had not been previously detected in sequenced cultures of s.s.. MM1 thus represents only the second s.s. strain to contain this plasmid and the first fully assembled. MM1 carries *vls* antigenic variation system on its lp28-8 plasmid which consists of the expression site *vlsE* and an array of 18 cassettes. The characterization of MM1 as a Midwestern USA isolate and a well-established s.s. strain in Lyme research and vaccine studies, contributes to understanding Borrelia diversity, and will facilitate the development of more specific diagnostics and vaccines.

## Acknowledgments

We thank Barbara Grimes† for initial culturing of the MM1 spirochetes used in this study and Dr. Qiang Tian for assistance with nanopore sequencing. We also thank Mary Brunkow for her guidance on the project and Max Robinson for his suggestions for the analysis. We gratefully acknowledge support from The Wilke Family Foundation, The Steven and Alexandra Cohen Foundation, Jeff and MacKenzie Bezos, and the National Institutes of Health, National Institute for General Medical Sciences, National Centers for Systems Biology Grant 2P50GM076547-06A1.

## Supporting Information

**S1 Fig. Coverage of MM1 sequence reads across *B. burgdorferi* B31 plasmids.** An initial mapping of MM1 reads to the type strain B31 suggested lack of plasmids lp21, lp28-1, lp28-2, lp38, lp56 and lp5 in MM1 genome, while the presence/absence status of other plasmids in MM1 remained ambiguous. B31 plasmids are shown in each plot with X-axis and Y-axis representing Reference Start Position and Coverage respectively.

**S2 Fig. Genome-scale dotplots of *Bb* MM1 and B31.** A) Dotplot of MM1 versus B31 shows a very high degree of similarity. MM1 contains a subset of the plasmids of B31. B) Dotplot of MM1 versus itself illustrates two characteristics of Borrelia genomes. First, the cp32 family of plasmids show great similarity. Second, a large repetitive region, the *vls* locus, appears as a prominent dark square on plasmid lp28-8.

**S3 Fig. Nanopore sequencing reads confirm the novel SNPs in *nifS* and *pepX***. Top: The *de novo* assembly of the *nifS* gene matches the MLST *nifS* type 12 except for a C->T polymorphism. Eleven nanopore reads align to this region and all contain the T variant. Bottom: The de novo assembly of the *pepX* gene matches the MLST *pepX* type 1 except for a T->C polymorphism. Nine nanopore reads align to this region and eight contain the C variant. The two SNPs of interest are noted with a red asterisk (*).

**S1 Table. MM1 gene members of PFam32, PFam49, PFam50 and PFam57/62 identified by spectral clustering of protein sequences**. The table lists the results of SCPS analysis for the 4 PFams 32, 49, 50 and 57/62 (see Materials and methods) in MM1, N40, JD1 and B31 strains. Genes located on replicons of the same type are arranged in nearby rows within each paralogous gene family. Dashes mark missing entries, which may fall below cut-off or may be missing in the respective strain.

## Footnotes

† Passed away June 16, 2017.

